# Recut: a Concurrent Framework for Sparse Reconstruction of Neuronal Morphology

**DOI:** 10.1101/2021.12.07.471686

**Authors:** Karl Marrett, Muye Zhu, Yuze Chi, Chris Choi, Zhe Chen, Hong-Wei Dong, Chang Sin Park, X. William Yang, Jason Cong

## Abstract

Advancement in modern neuroscience is bottlenecked by neural reconstruction, a process that extracts 3D neuron morphology (typically in tree structures) from image volumes at the scale of hundreds of GBs. We introduce Recut, an automated and accelerated neural reconstruction pipeline, which provides a unified, and domain specific sparse data representation with 79× reduction in the memory footprint. Recut’s reconstruction can process 111 Kneurons/day or 79 TB/day on a 24-core workstation, placing the throughput bottleneck back on microscopic imaging time. Recut allows the full brain of a mouse to be processed in memory on a single server, at 89.5× higher throughput over existing I/O-bounded methods. Recut is also the first fully parallelized end-to-end automated reconstruction pipeline for light microscopy, yielding tree morphologies closer to ground truth than the state-of-the-art while removing involved manual steps and disk I/O overheads. We also optimized pipeline stages to linear algorithmic complexity for scalability in dense settings and allow the most timing-critical stages to optionally run on accelerated hardware.

## I. Introduction

Morphology, the 3D shape of single neurons, and topology, the arrangement, coverage and connectivity of such cells, may be a contributing facet of neuronal function and pathology. Morphology in particular is a critical component of cell type taxonomy: the process of discriminating and classifying different neuron types by genetic, proteomic, connectivity or pathological characteristics[1]. A common method to studying morphology is reconstruction, the process where fluorescently labeled cells in 3D images are segmented and compacted into ball and stick models[2]. Recent reviews have identified a scalable[3], end-to-end[4] pipeline tool for single-cell reconstruction as the greatest need of the community. The Recut pipeline comprehensively addresses these needs.

Reconstruction establishes a *coverage topology* of the segmented neuronal regions which can be alternatively represented as a set of vertices with connections i.e. a graph. This graph *G* = (*V*, *E*) is composed of vertices *v_i,j,k_* and edges *e*, where each vertex has a fixed and unique location in 3D space. The neuroscience community and reconstruction tooling has aligned on the SWC standard[2], a text file format where each line specifies a vertex in the reconstructed graph with its 3D coordinates, radii and parent vertex. Since vertices only carry a directed edge to their parent, the SWC format actually describes a *tree* as opposed to a general graph. Once the tree of all neurons is constructed, it is partitioned into subtrees that represent individual neurons. With individual neurons established, their morphology is analyzed in aggregate.

### A. Challenges and Related Work

#### 1) Accuracy

Dozens of computer-assisted implementations exist for reconstruction as documented by comprehensive literature surveys and reviews[5], [4]. Graph-based reconstruction methods are particularly suited in modeling morphology and connectomic data and are deeply embedded in the analysis patterns of the neuroscience domain. However, they are prone to suffer accuracy loss from a variety of factors such as choice of background threshold value, normalization of the image, and pixel-level noise, all of which contribute to erroneous path breaks.

Due to incorporating larger spatial context cues at various window sizes and resolutions and training, neural network (NN) methods are a natural complement to mitigate the issues of graph methods. While this removes sensitivity to free parameters such as background threshold, it requires task-specific model selection, a robust training set and still must handle the conversion to the tree-like SWC format for downstream algorithms such as in Figure 1.

**Fig. 1.**
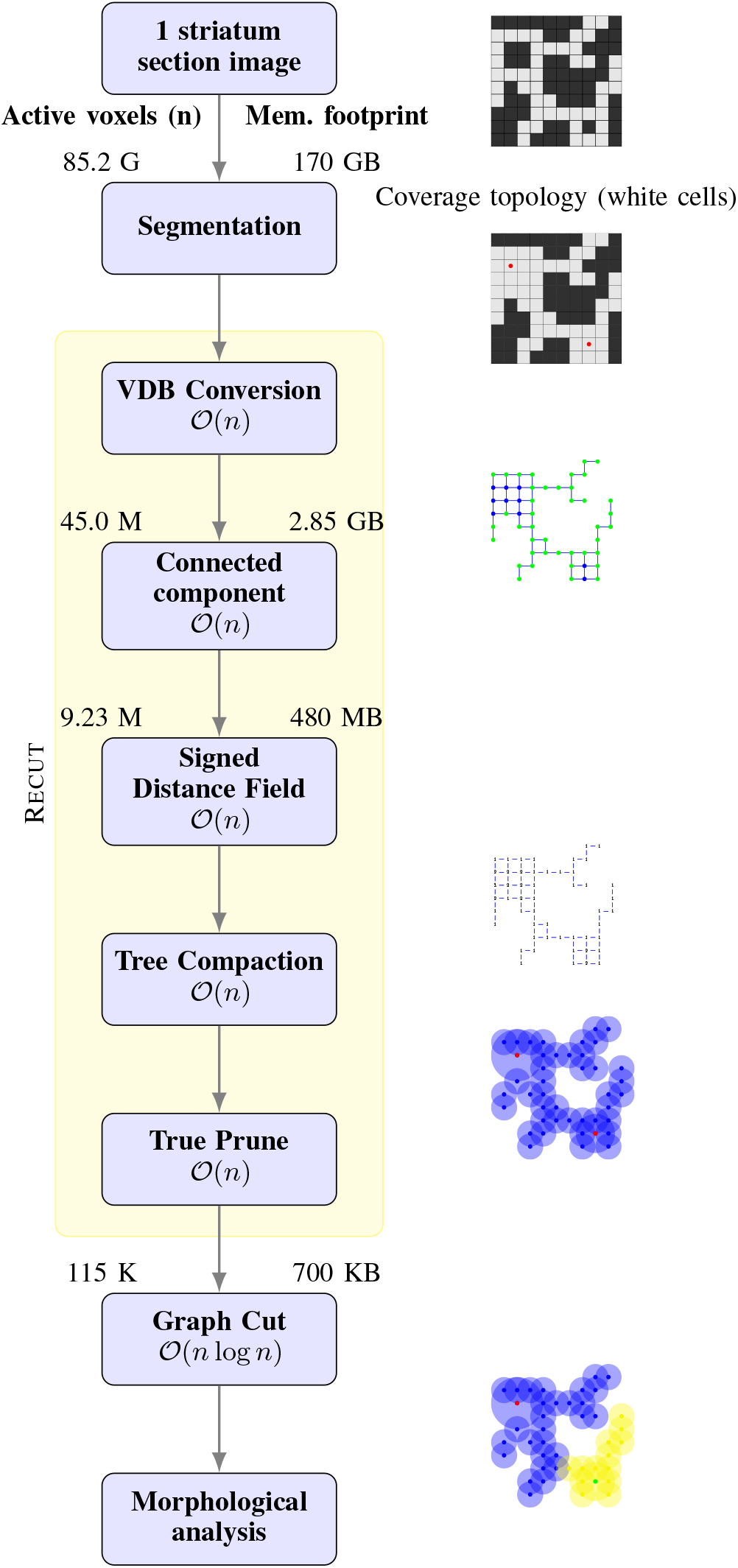
Reconstruction progressively compacts the data footprint across the pipeline stages: starting with 3D images, we convert the representation into a sparse graph then further partition into individual graphs for each neuron, then finally aggregate for morphological analysis. With compaction, Recut’s throughput is limited by the DRAM bandwidth of a system. Therefore the uncompressed footprint in memory of the active voxel working set (n) is shown on the right at each narrowing. Numbers shown are for a 30x objective lens non-downsampled section of the striatum brain region.

In a recent 27 method comparison[6], all top mouse algorithms were graph-based[7][8][9][10], with the exception of Advantra[11] which uses the Monte Carlo method at the initial stage. The top performing software Neutube[7] as well as many off the shelf reconstruction tools [12] employ a fastmarching algorithm similar to APP2. The connected component (CC), signed distance fields (SDF), and tree compaction (TC) stages of this paper therefore compare to the algorithms in APP2 as baselines for performance accuracy metrics.

#### 2) Algorithmic Efficiency

Existing graph based reconstruction methods have comparatively low data access and computation. In conventional high-resolution light microscopy methods such as confocal (see details of our imaging pipeline in section 2.7), the *coverage topology* foreground is a tiny proportion of the total image. In such situations, graph approaches can avoid excessive computation since they avoid the dense and redundant access patterns found, for example, in convolution. Graph methods are therefore a strategic starting point to base further optimization effort upon. We refer to the metric of computation with respect to data size (e.g., image voxel count or *n*) as *algorithmic efficiency*. We compare algorithms based on their theoretic computation counts expressed in terms of *n* disregarding constant factors based on the big 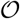 notation[13]. This notation is fundamental in understanding and comparing efficiency. For instance, in the same comparison study[6], APP2 had the best runtime performance due to utilizing the fastmarching algorithm. Yet even APP2’s radius calculation and pruning stages have sub-optimal algorithmic efficiency which severely limits the data sizes they can feasibly run on.

#### 3) Performance and Resource Utilization

Better algorithm choice is an entry point for faster software, in practice, however hardware-specific optimizations yield far greater speedup factors[14]. Neuroscience pipelines demand unprecedented scale exposing a long-standing weakness in computing infrastructure: data movement. Graph methods often traverse image volumes of sizes beyond memory capacity. Smaller subregions are then streamed as needed at runtime to mitigate large reads. These subregions are often visited and therefore read repeatedly and a large fraction of these dense regions are background or unaccessed pixels. This equates to programs that spend several orders of magnitude more time reading background values from disk than performing the desired computation. Additionally, generic compression methods on disk do not leverage the inherent spatial sparsity of neuroscience data.

Algorithms are tightly coupled with data structures [15] and a unified vertex and data representation for all algorithms is essential to prevent excessive movement related to data conversions and inefficient access patterns. An ideal data model supports all desired algorithms efficiently and is leveraged as early as possible in a multi-stage pipeline. We choose the Volumetric Dynamic B+ tree (VDB) format [16] since it is optimized for a fast access and low memory footprint for spatially sparse volumetric data. VDBs have a hierarchy of cubic regions for exploiting sparsity at various granularities. When a subregion contains no active voxels or entirely uniform values, its representation can be collapsed into a single value as visualized in Figure 2.

**Fig. 2.**
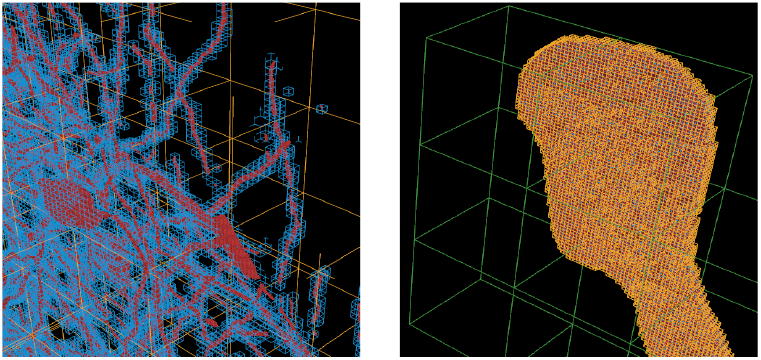
VDB grid of sparse neuronal data. The coverage topology of all active voxels is shown in red. Active leaf node regions–those that contain an active voxel–are outlined in blue. Active internal nodes regions are outlined in yellow and active secondary internal nodes in green. Empty leaf nodes and internal nodes have no outlines and have tiny memory footprint overheads.

#### 4) Scalability

Neuromorphology literature cites reconstruction as the bottleneck[17], [4] for high-scale neuroscience. Graph kernels such as pruning are also cited in neuroscience research as the bottleneck to analysis[6]. Graph reconstructions methods lack *scalability*–the ability to handle large image regions on common systems such as a laptop or even on modern servers. In practice, this is due to the combination of the efficiency and performance concerns discussed above. Adapting to modern hardware requires enabling concurrency throughout a pipeline which can mitigate such data concerns. APP2, like other available iterative reconstruction methods, is only single-threaded. This is why large-scale applications must start multiple APP2 programs for each soma to enable some concurrency [18]. Poor scalability is such a concern that common practice involves *windowing* regions of an image which can create cutoffs, artifacts, and intensive human intervention.

#### 5) Automation

Large barriers to automation in reconstruction still exist. Even trivial manual steps vastly lower throughput by creating forced stops and long down times which greatly degrade the value of performance improvements. Lack of algorithmic efficiency or scalable design itself creates new barriers to automation (e.g., enforcing windows of neurons to be cropped by hand which takes about 2-3 days per brain even when accelerated by an automated cropping program). Additionally, SWC outputs of semi-automated tools are usually filtered by hand. Other semi-automated pipelines such as [18] must manually identify soma locations since they are so critical to accuracy, but requiring intervention so early can limit many benefits of automated reconstruction. Finally, reconstruction is sensitive to signal-to-noise of the reporter signal, which can be altered by tissue preparation, imaging, and stitching.

#### 6) Correctness

Vaa3D and the BigNeuron project have significantly advanced collaboration in the field by providing a framework to compare reconstruction method accuracies. However, reporting the end-to-end accuracy of reconstruction paper is only a first step towards reproducible research and reusable software tool-chains. For long-standing software that is intended to grow with collaborative effort, individual components of functionality also need to have corresponding automated unit tests. In order to improve algorithms and modules, baseline behavior must be first quantified and verified with such tests[19].

#### 7) Portability

The neural anatomy and morphology community has had inspiring success in aligning data formats (e.g., with the tree-based SWC format) and registration atlases (e.g., CCF). However coordinating software development collaboration is far more complex but has even greater potential benefits for sharing extensible reconstruction, analysis, and visualization pipelines.

Software is far more dynamic, usually with a complex set of shifting dependencies that aren’t declared explicitly by authors. The implications towards reproducible results are a growing concern, and other large scale scientific projects such as at CERN have adopted sophisticated methods to robustly lock and archive analysis software and all of its dependencies[20].

### B. Contributions

To evaluate our implementation we provide several features and principles:

- *Correctness*. Recut’s automation or computational techniques produce results consistent with the original intent without unexpected or unknown behavior changes or failures.
- *Accuracy*. Our framework demonstrates comparable or higher accuracy to those established in the reconstruction community.
- *Algorithmic Efficiency*. Recut demonstrates high algorithmic efficiency by reducing unnecessary computation and data access.
- *Performance and Resource Utilization*. Recut’s implementation includes hardware specific optimizations to improve metrics like latency and throughput in ways that are as portable as possible.
- *Scalability*. Our software design is scalable, streamable and concurrent, meaning the implementation natively supports and performs consistently when adding significantly larger data sizes or more processors.
- *Automation:* Recut automates soma detection and first pass reconstruction and single-neuron partitioning from clusters. Manual proofreading steps are pushed as far downstream in the pipeline as possible to enable the benefits above.
- *Portability*. Open source software implementations are increasingly critical to scientific endeavors but have complex dependency chains that can take even software experts days to set up and integrate properly. Recut is installable and usable as a standalone program or a C++ library with 2 commands from a Unix command line.

## II. Pipeline

We refer to the provided algorithms as *stages* since they must proceed in a specific forward order for a given image region. Early stages of the pipeline operate on dense regions, whereas downstream stages tend to operate on an abstracted and sparse tree representation. The CC, SDF, and TC stages of Recut have a direct mapping with algorithms or components of APP2 and the intended transformation semantics are matched as best as possible. However, since the implementations are different we use the standard names of the transformations as they are used in the literature.

### A. Image Preprocessing (IP)

The pipeline takes as input stitched image volumes. Inputs can be further enhanced to improve the fidelity however for discussions in this work we simply segment the raw inputs.

### B. Segmentation (SG)

To mitigate issues of raw images and membrane labeling, our pipeline applies the artificial neural network based algorithm U-net[21] to segment all regions of the image belonging to neural tissue. This segmentation stage outputs a binarized dense volume the same dimensions of the original image with voxels that label neurons.

Additionally, this inference step aggregates a list of the filled soma locations. Soma location is critical for downstream stages since it acts as a root point for the branch-like projections–*neurites*–characteristic of neuron morphology. Somas are used to establish the starting seed locations for the subsequent stages of the pipeline. Somas are generally visible in image space and so critical to reconstruction accuracy that many pipelines rely on manual human identification of these locations. However, we rely solely on our automated methods which is essential for exploiting parallelism and boosting throughput.

### C. VDB Conversion (VC)

After segmentation, we convert the dense NN outputs into the low memory footprint VDB grid. This step conducts a dense read 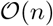 such that all later stages of the pipeline have fast accesses on the sparse volumetric grid.

### D. Connected Component (CC)

Segmentation establishes contiguous regions of coverage topology based on contextual information in local regions of the image. However, only a subset of these labeled regions are biologically plausible. In order to be relevant, any labeled voxel must also be reachable from the determined soma locations. Given the VDB graph with filled and identified somas as starting locations, the connected component stage labels all reachable vertices as selected in a region growing scheme. This stage yields all components of the image and the encompassed somas. *Component* refers to a cluster of neurons that are connected by at least 1 voxel. This CC stage fills a similar purpose to APP2’s fastmarching stage or other reconstruction algorithms that use a Dijkstra or single source shortest path algorithm (SSSP) to traverse foreground voxels. Contrary to these other algorithms, our CC stage simply stores a parent for each selected vertex with no distance or salience field.

### E. Signed Distance Field (SDF)

After the CC stage, the working set–all active elements–is complete. In other words, all voxels belonging to the coverage topology are known. However, the CC stage selects a vertex for each foreground pixel, which is highly inefficient for subsequent stages in the pipeline. To compact the redundancy in selected vertices, each vertex is considered as a 3D sphere with a radius that extends to the nearest background pixel (inactive voxel). Calculating these distances provides a coverage area of each vertex via an operation referred to as SDF.

### F. Tree Compaction (TC)

Given a set of selected vertices with radii, we can perform graph compaction by starting from somas or more salient nodes first and traversing the graph, removing vertices that are already within the covered radius of a previously visited vertex. This leaves a graph that covers the same active image volume but is described by far fewer vertices. In neural datasets, this reduces vertex counts as detailed in Table VI.

### G. Tree Prune (TP)

While it is possible to compact losslessly for an exact volume coverage of foreground, in practice this is not useful since it produces enormous numbers of spurious branches that obfuscate morphology metrics. These adjustments traverse the active set of vertices and apply refinements on the sparse graph. In our pipeline, we use TP to remove all branches of length 1 with a parent bifurcation point.

### H. Graph Cut (GC)

Recut implements a method to separate connected components that contain multiple somas (graphs) into individual trees (SWCs) based off the parent connection assignment in the CC and TP stage. This allows Recut to remain a practical end-to-end solution in dense labeling settings. For more accurate partitionings of the graph in even denser settings, the user can optionally run the method offered in [22].

### I. Windowing, Visualization and Proofreading

Recut compresses the full image bounding volume to the VDB format then reconstructs and outputs discrete SWCs with their exact corresponding bounding window. This window is reflated from the foreground VDB voxels and written to a TIFF image on disk for subsequent visualization during proofreading in common morphological GUI software such as Neutube or Vaa3D. Recut can optionally output SWCs or point VDBs with coordinates embedded in the original image for correcting breaks where the path extends beyond the components bounding volume. This can be particularly useful for visualization in software tools such as Houdini that natively support both the VDB image and VDB point grids output by Recut.

### J. Morphological Analysis

The SG and CC stages entirely determine the coverage topology accuracy of a reconstruction in our pipeline since subsequent stages merely compact the graph representation. However, current analysis techniques focus on graph morphology which is more abstracted than the volumetric and position information afforded by coverage topology. *Persistence homology* for graph similarity builds a feature space that can express the morphological measurements of Sholl analysis while retaining spatial embedding information[23][24]. Persistence images have been used to classify neuronal types with subtle morphological differences that are difficult for even human experts to discriminate [25]. We leverage the Topological Morphology Descriptor (TMD)[23] an implementation of persistence homology. TMD is less sensitive to imaging and the accuracy of the SG stage than pixel based accuracy metrics such as those used in [6]. The TMD feature space also has particular bearing on the edits to a tree that occur during proofread such as fixing breaks, merging branches, etc.

## III. Implementation

### A. Algorithmic Efficiency

As demontrated in Figure 1, we have two basic types of algorithms: NN and graph-based. NN is currently only used for the SG stage. Whereas graph algorithms tend to have sparse irregular data access with complex concurrency schemes, NN methods have dense and regular data access patterns and simple parallelization schemes. The graph methods are also particularly well suited to take advantage of sparsity.

The desired transformations are quite common and there are many alternate implementations to these stages such as pruning with sphere packing, however CC and SDF stages are graph-based employing a breadth-first search on an iterative, advancing wavefront. This allows us to use the same underlying graph representation, overhead data structures and traversal patterns, thus greatly simplifying the implementation and reducing memory communication. The efficiency of the graph stages of Recut are listed in Table I.

**TABLE I.** A comparison of the algorithmic efficiency of Recut not accounting for parallelism over multiple processor cores. *n* indicates the total working set count of vertices at that particular stage, *r* indicates the radius distance of a particular node.

| Stage | APP2 | Recut |
| --- | --- | --- |
| CC | - | $\mathcal{O}(n)$ |
| FM | $\mathcal{O}(n \log n)$ | $\mathcal{O}(n \log n)$ |
| SDF | $\mathcal{O}(nr^3)$ | $\mathcal{O}(n)$ |
| TC | $\mathcal{O}(nr^3)$ | $\mathcal{O}(nr^3)$ |
| TP | $\mathcal{O}(n)$ | $\mathcal{O}(n)$ |

Besides TC, GC, or the optional fastmarching, the algorithmic efficiency of the graph stages of Recut are 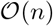 as depicted in Table I. This is ideal efficiency since each stage needs to update *n* nodes and those updates are not dependent on the order of *n*.

The FM algorithm is less efficient than Recut’s CC stage because it requires on the order of log *n* more work per visited node to run. Note that we include the term *r*–a node’s radiu–seven though it is independent of the working set count *n* since it helps in understanding the performance of the stages below. APP2’s calculation of radius SDF algorithm requires *r*^3^ operations per node, since it checks a circle, *r* times, whereas Recut’s implementation is again on the order of 1 operation per node, due to the same implementation pattern as the previous FM stage. Recut’s TC stage sparsely checks a sphere of vertices according to each node’s radius, whereas the APP2’s hierarchical prune densely checks a circle, *r* times. TP for both Recut and APP2 require a single traversal of the compacted graph.

### B. Resource Utilization

#### 1) Vertex Representation

We utilize a customized vertex model that can support downstream stages while remaining compact as shown in Tables II and III. Neuroimaging data has inherent geometric properties and those properties tend towards predictable power law distributions. Upon statistical analysis we can enforce data value range cutoffs to quantize the types of fields used at each vertex to constrain the vertex memory size. Recut employs two basic types to represent the finest granularity of an image: the *vertex* and the *message vertex*. Any updates occur at the granularity of a vertex. All vertex attributes are held within VDB grids, whereas message vertices are temporary and are emplaced dynamically in each FIFO. Concurrency is achieved via messages between leaf nodes which are buffered in FIFOs. Message vertices are simply an extension of the vertex type with a slightly larger memory footprint since they must be tagged with an offset to distinguish their coordinate identity within a leaf.

**TABLE II.**
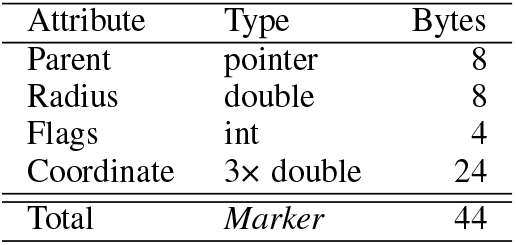
Vaa3D, APP2 marker footprint in memory

**TABLE III.** Recut *vertex* and *message vertex* footprint in memory

| Attribute | Type | Bytes | Red. v. APP2 |
| --- | --- | --- | --- |
| Parent offset | 3× int8 | 3 | 2.7× |
| Radius | uint8 | 1 | 8× |
| Flags | uint8 | 1 | 4× |
| Message offset | 3× uint8 | 3 | 8× |
| Total | Vertex | 5 | 8.8× |
| Total | Message Vertex | 8 | 5.5× |

#### 2) Image Representation

Typically, acceleration or parallelization schemes of graph processing rely on a preprocessing step to partition or arrange nodes and edges according to connections. This is an effective data-oriented approach for kernels operating on static power-law graphs[26]. Unlike most high performance graph kernels, the graph stages of Recut are dynamic: they prune and reassign parents (edges) and repeatedly prune substantial portions of the active vertex set. Due to their embedding in volumetric space, the graphs are also planar, with an edge count of 1, with highly local connections. Additionally, only a small fraction (demonstrated in Table VI) of all image voxels are ever visited and selected during the CC stage. For these reasons, the common high performance graph arrangement schemes would mostly be single-use requiring a costly rebuilding during or after each stage. This rearrangement itself would have a similar runtime as a stage.

To deal with these issues, Recut’s internal image and graph format both use OpenVDB grids, which are optimized for fast access and low disk and memory usage for sparse volumetric data[16]. VDB grids allow Recut to transition from an image abstraction to a graph representation seemlessly, which is why we use the terms voxels and vertices interchangeably. VDB grids hierarchically partition the data at different flexibly-sized cube regions. The smallest partition region in a VDB grid is termed a leaf node, which is a cube of vertices processed by a thread independently of other leaf nodes. The leaf node size introduces spatial and temporal locality by enforcing a thread to stay within a small region of data until it completes processing all possible updates. Vertex attributes are accessed or stored as needed at the granularity of leafs. Each active leaf node requires a FIFO to hold future vertices to process and receive incoming messages.

#### 3) Coverage Topology

Recut is designed to operate on arbitrarily large images that represent contiguous volumes of brain space. For example, light sheet data produces whole brains which Recut ingests as a single image, efficiently combines or compares multiple channels, and translates to a single topology representation. We conceptually split the topology into two parts, *coverage* and *attribute*, which corresponds in concept to the underlying OpenVDB data structure’s usage of bit masks and attribute arrays respectively. The SG stage produces an overrepresentation of the topology *coverage*–the total segmented voxels of the original image–which the CC stage constrains to only vertices reachable from somas. After the CC stage, the coverage is complete and the voxel based accuracy of the reconstruction is established (although paths can be reassigned to different neurons during GC). While *coverage* may be static after the CC stage, the *attribute* topology–the working set of all active vertices and their properties–gets reduced during TC. Despite the narrowing in memory footprint, the intended coverage is preserved abstractly via the radius property of the remaining vertices in the attribute topology.

#### 4) Overhead

Recut allocates a FIFO for each of the active leafs determined by SG before running the CC stage. The subset of these FIFOs, that are reachable from somas, have a lifetime for the rest of the pipeline since they need to dynamically buffer vertex messages to any leaf to ensure thread safety. We reduce memory requirements needed for message buffers further by leveraging an update grid which sparsely stores whether border regions have been updated such that messages can be generated safely in the integration method at the end of each iteration shown in Algorithm 1.

#### 5) Memory Leakage

Mismanagement of the dynamic memory footprint–*memory leakage*–leads to poor utilization of available resources and could cause large drops in program performance (i.e., scalability issues). More subtle issues in design can lead to memory leakage. For example, while a raw image buffer is used for segmentation, keeping it in program memory for the remainder of the stages would severly limit the regions that could be processed in parallel. This leads to a fundamental design point, only the minimum data needed for computation is accessed at each stage. Deep pipelines with diverse algorithmic needs require tradeoffs between the memory footprint, the algorithmic expressivity and the fast access. For example, Recut builds a tree graph before the GC stage by enforcing that each vertex has 1 edge: its parent. If Recut were expanded to include algorithms requiring undirected graphs such as GC, it would substantially increase the memory footprint of a vertex as it would additionally need to hold antiparallel edges (its children).

#### 6) Sparse Adaptability

While the above details current design choices for our pipeline, almost all aspects can be tailored to other needs. Fluorescent labeling sparsity can vary greatly but is generally fixed across many imaging rounds. The sparsity displayed in neuromorphology is not a random set of pixels, it has localized correlation and spatially identifiable coherence, a property known as *spatial sparsity* [27]. Furthermore, exploitable spatial sparsity tends to increase with higher dimensionality, resolution, expansion factors or added imaging channels.

The expected labeling density can vary between 0.001% to 0.1% in order to exploit ultra-sparse scenarios. Recut can avoid treating the region as dense and instead monitor the sparse set of active vertices in a data structure optimized for low memory footprint and fast access for sparse volumes known as OpenVDB[16]. User-specified compile-time constants control this varying behavior and linkage with the optional OpenVDB 3rd party library so that runtime latencies are avoided.

#### 7) Downsampling Flexibility

It is common in visualization and graphics to encode images at various resolutions–full resolution, half resolution, quarter resolution, etc. This technique is used in high performance applications and the HDF5 format, and is most commonly referred to as multi-resolution or mip-mapping. This method can reduce the speed of accesses in high performance applications at the cost of doubling the memory footprint. The VDB library provides a method to build multi-resolution sparse grids, but the sophistication of VDB data structure allows far faster access and smaller footprints without needing to adapt algorithms to this paradigm. While the resolution of voxels in grids is fixed throughout Recut, it would be possible to simulate upsampling by interpolation or downsampling by access or traversal pattern at runtime using VDB’s provided functionalities.

#### 8) Memory adaptability

Depending on the spatial sparsity characteristics, VDB grids can be arbitrarily deep with adaptive grid sizes chosen specifically for a data sample or type. This can be leveraged to substantially reduce the memory footprint at the cost of slower access times. To explore this tradeoff further refer to [16].

#### 9) In-Memory Computing

Combining sparse data structures and a minimal vertex representation allows substantial reductions in memory footprint as detailed in Table VI. These reductions can shift the performance bottleneck from being disk-bound to main-memory bound (DRAM) for our entire end-to-end pipeline on a reasonably equipped imaging workstation. System builders of imaging workstations for sparse data should therefore keep in mind that in-memory computing will have disproportionately large benefits for performance with Recut.

### C. Scalability

Due to the low reliance on global memory structures, we designed a lock-free shared memory approach based on [28] that operates on VDB grids that represent the required graph data. The basic structure of this scheme is illustrated in Algorithm 1. Concurrency is foremost at the granularity of leaf nodes via the Intel TBB library, which parallelizes leaf processing across threads on a single CPU. We evaluate this framework via the speedup factor with respect to the number of cores used. Performance of these kernels is highly data dependent, therefore we define scalability metrics in terms of cores since they are aligned and localized with the CPU cache hierarchy more than threads.

Recut can traverse through VDB grids following neurite paths in 3D space. However, since vertex attributes are stored in distinct regions of memory (in struct-of-arrays fashion), only the attributes of the active leafs specifically needed by each stage are accessed.

The VDB implementation supports image regions with x,y,z coordinates between −2e9 to 2e9. Full adult mouse brain confocal images imaged with a 30× objective lens have a bounding volume of about 65k×28k×2.7k with anisotropic z downsampling. A single VDB grid bounding volume can encompass 1.8 e16 of such brains arranged side by side due to the reliance on indexing via 3-dimensional coordinates of type signed integer-32. VDB grids are specifically designed to store the spatial structure in volumetric data and compress uniform or background regions. Our data has complex spatial sparsity at multiple resolutions which would not be effectively exploited by more naive adaptive grid methods. We evaluate the cost of the VDB format via the footprint in memory with respect to the active voxel count of the labeled segmented regions. This metric should scale linearly for suitable sparse data structures.

### D. Automation

The segmentation stage automatically identifies soma locations and removes the need to specify a background threshold value or do other image processing. Alternatively, Recut can take raw image buffers and a desired foreground percent. While a desired percent is less fraught than specifying a background intensity value as with APP2, it can still be a trial and error process which is why the automated SG stage is recommended.

#### Algorithm 1: This pseudocode represents the critical update loop. Each graph processing stage executes this loop with a stage specific traversal and a vertex field contained within the function on line 8. Integrating the ghost cell updates contains the added overhead of the parallel algorithm. Both inner parallelized for loops implicitly induce synchronization overheads.

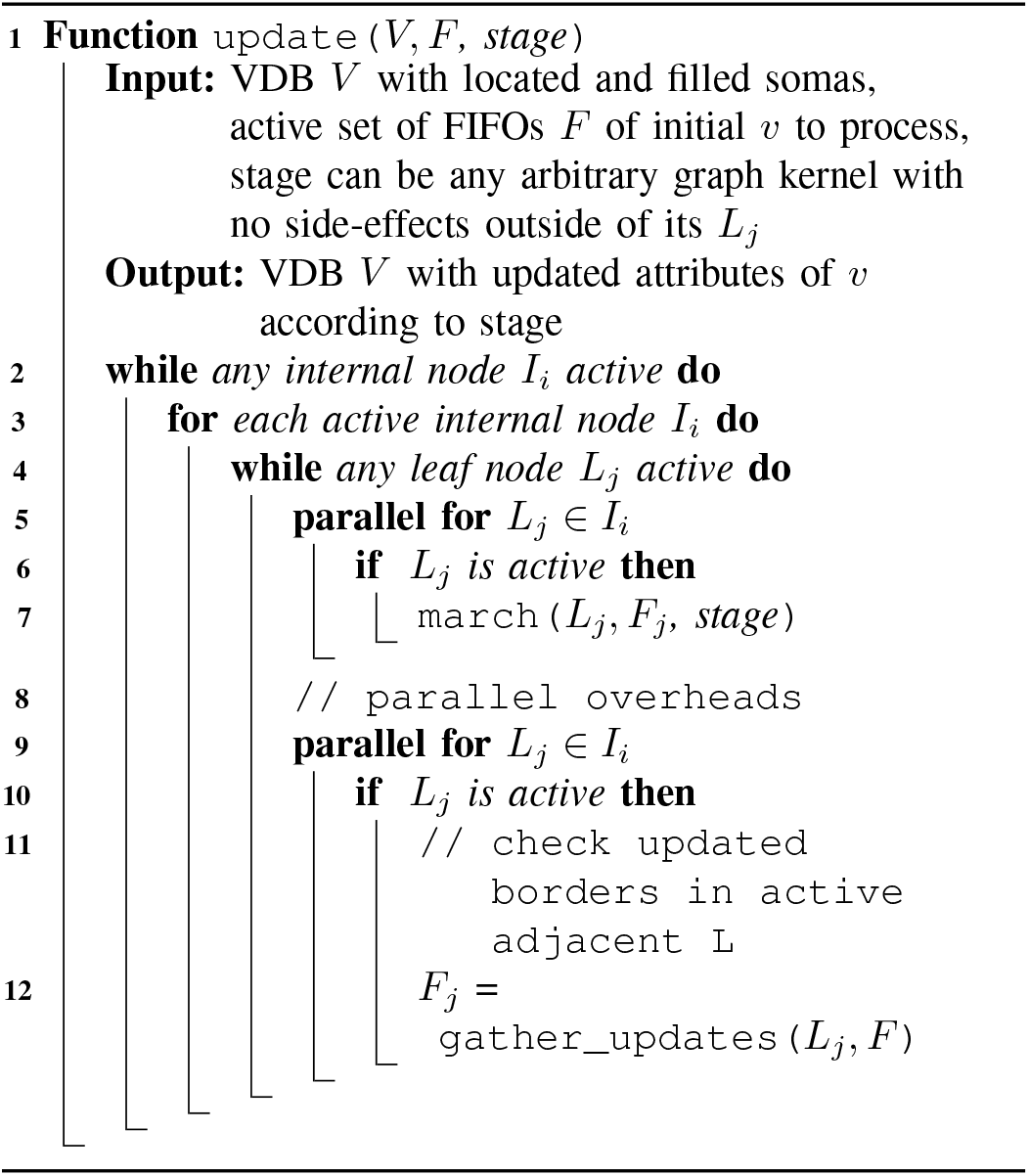

### E. Correctness

To ensure basic correctness, modern application development handles implementation complexity by *test-driven development*. This is a paradigm whereby a program’s isolated functionalities or integrated behavior is verified by a test suite accompanying the software. Rather than rely on developers to run these tests by hand to ensure no bugs have been introduced, full test suites can be run automatically on code changes or on continuous schedules to prevent any correctness regressions. This automated process, known as *continuous integration* (CI), enables large teams to deploy changes to complex software systems rapidly with confidence. Recut was developed in conjunction with a set of fast tests (less than 2 s total runtime) that are run automatically by the hosting platform.

### F. Portability

At this initial release, the Recut framework is written in modern C++ (C++17 and above). It leverages language supported threading, atomic variables and some lightweight third party libraries. Recut optionally links with different image reading software as image data types are varied and nonstandardized in the neuroscience community. In this paper, we link with an internal library to read TIF or Imaris (HDF5) images. Additionally, we leverage python functionality for the segmentation and graph-cut step detailed in [22]. Recut and all its dependencies are distributed via the Nix package manager such that installation, compilation, and development requires running only two commands on a Mac, linux, or PC with WSL installed.

#### Algorithm 2: Connected component → 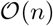

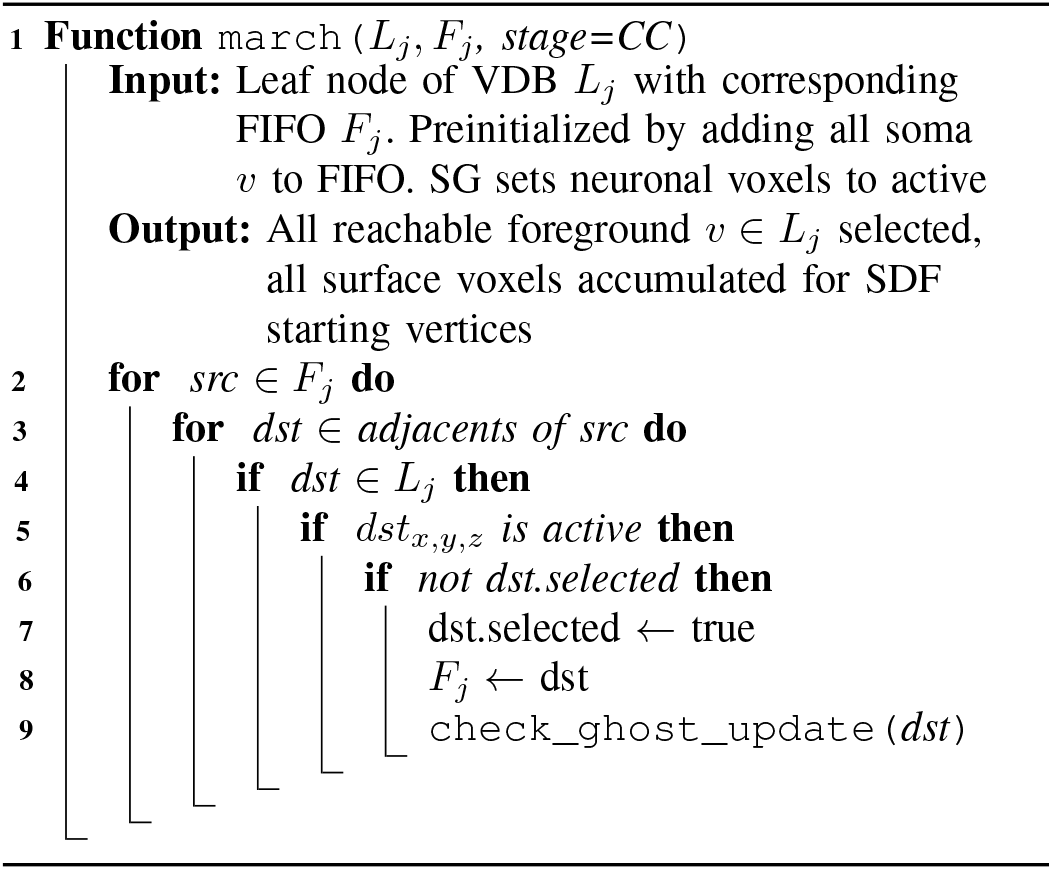

#### Algorithm 3: Signed Distance Field → 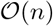

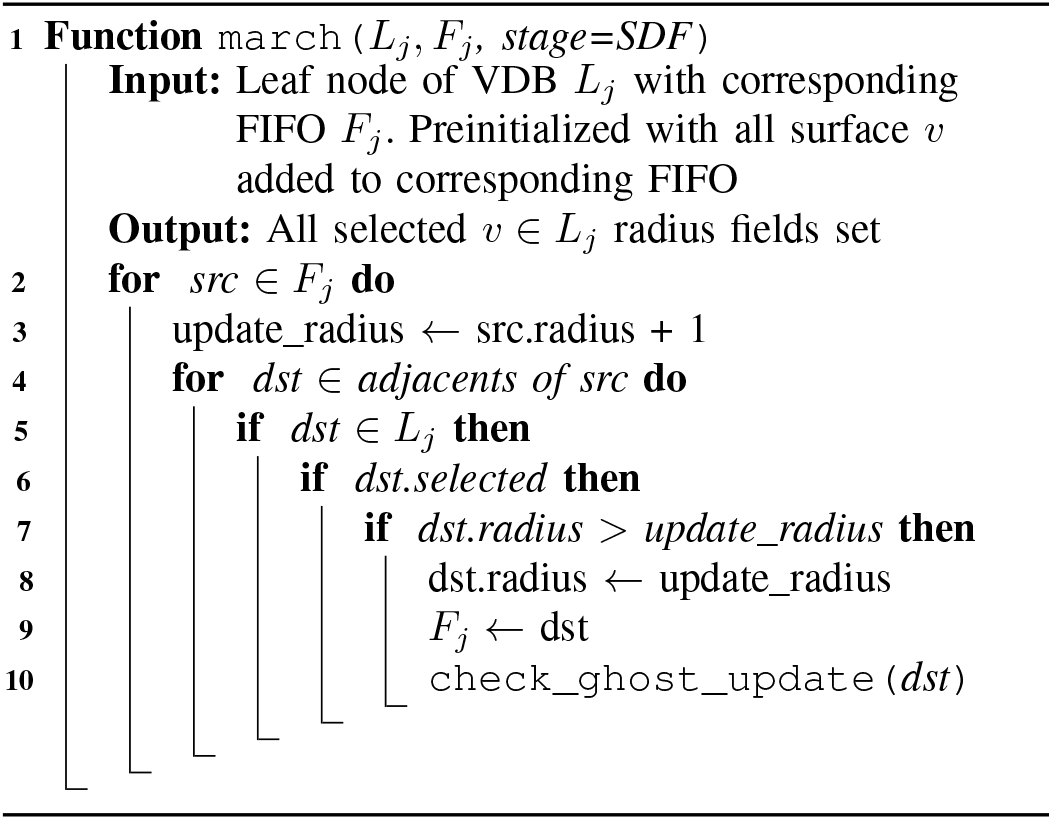

## IV. Experiments and Results

### A. Data and Setup

This pipeline takes TIF files as single-channel gray scale 3D images at 16-bits per pixel. The gray scale intensity represents the fluorescent labeling density determined by wet lab fluorescent techniques. The measured labeled signal density % is detailed in Table VI. Labeling is a product of months or years of biochemical tuning and research efforts which vary by scientific problem. If a fluorescent label is designed with too low a labeling frequency, the yield of neurons per mouse brain or breeding population may be too low to produce statistically viable results. If a fluorescent labeling is too high, neurons become a mesh, erroneously connected from the perspective of the reconstruction technique.

The neural data used in this report is membrane-labeled mouse brain data with about 0.05% pixel label density determined at the SG stage. For real data testing we use a 7680×8448×383 image (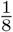 coronal section; 2.36e10 total pixels) of mouse cortex labeled using CAM-K2 knockout at full resolution. The z dimension is 5× smaller than *x* and *y* (covering steps of 1 um vs. 2 um respectively). This region contains 232 distributed somas with dense mesh-like, continuous branching patterns. 1 section is about 16× the window size allowed by most reconstruction software.

We tested the performance on a workstation with an AMD Ryzen Threadripper 3960X 24-Core CPU at 3.79 GHz with 2 hyper-threads per core, 256 GB DRAM, and 3× Samsung SSD 860 EVO 4 TB configured as RAID0.

### B. Accuracy

At this labeling density reported, APP2 can only reliably terminate on data sizes up to bounding volumes of about 2048×2048×512 or about 2 gigavoxels before runtimes become excessive even on a modern CPU as noted in [6] and verified in our test suite. At 30× objective lens with our particular labeling probe, this bounding volume can encompass about 1 neuron’s span as shown in Figure 4. All accuracy comparisons to APP2 are for single neurons.

**Fig. 3.**
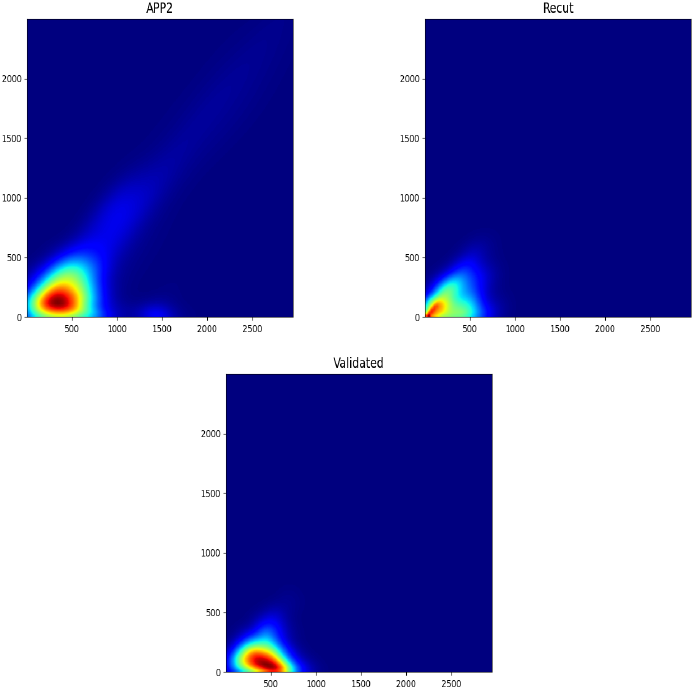
Persistence images for ground truth, Recut and APP2. Persistence images are aggregates of persistence histograms[23] a value for each branch is plotted by the radial distance from the soma at its birth (terminal) on the x-axis, to its death (bifurcation point or soma) on the y axis.

**Fig. 4.**
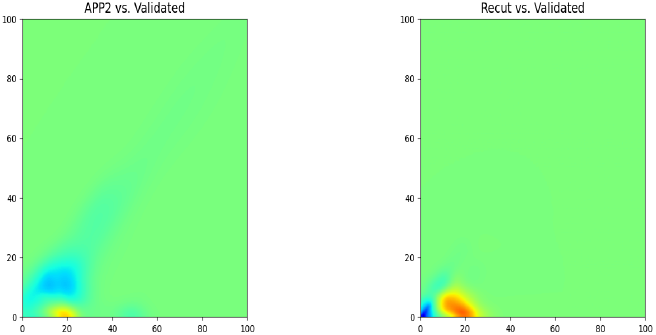
Recut or APP2 average persistence images subtracted from ground truth. Positive normalized values in red, negative in blue.

Our parallel connected component and SDF stage converge to exact equivalence with sequential runs and ground truth over synthetic test and real neural data as expected and confirmed by unit tests. Note that Recut’s exact matching with APP2 sequential results diverges from the SDF stage onward since APP2 implements a 2D simplification of distance to compensate for the slow performance of its algorithm. This APP2 behavior creates an error rate that increases with pixel size. We found for the scales of a soma (20-30 voxels) APP2’s error rate reached up to 23.1%.

APP2 employs hiercharchical pruning, a heuristic strategy to encourage the selection of a pruned set of maximally covering minimally redundant final vertices whereas Recut employs an optimized compaction strategy based off of an optimized VDB implementation of the Advantra prune method[11].

To compare the different approaches, we compare the absolute difference in average persistence images between APP2 and proofread ground truth (205) and Recut and validated (80.6) for N=42 sample windows with a single neuron. Note that the set of proofread validated samples were derived directly from the APP2 generated SWC. Thus, the difference indicates manual morphological edits that humans made. Recut’s lower difference scores indicate that the morphological characteristics are closer to final proofread ground truth.

The persistence images were generated from path distance thus the neurite diameter information is not included in these statistics. The ground truth persistence image shows that the majority of branches start at a similar distance from the soma (x-axis) and terminate even closer to the soma (y-axis) which corroborates the bush-like morphology of D1/D2 neurons. Persistence images for APP2 have higher values along the diagonal y=x which indicates that it introduces erroneous short branches beginning and ending at similar distances from the soma. The left-shifted peak of the APP2 compared to the ground-truth possibly indicates that long paths are either added or extending during proofreading. Additionally, the image for Recut suggests that it often fails by introducing unnecessary branches. These branches are short and particularly common closer to the soma. Although the recut image does seem to capture some of the peak of the validated ground truth, the image is dominated by small branches.

Note that Recut’s CC stage can be optionally run using a priority queue (heap) changing the stage to a parallel fast marching (FM) algorithm. Choice of FM vs. CC effects the CC traversal, the final parent paths back to root and the compaction order of TC. However, we did not find significant effects on the final accuracy by using an FM approach; thus, we omit it from the end to end accuracy comparison. However, researchers should compare and evaluate reconstruction algorithms via the respective effects on the final biological metrics they study.

### C. Algorithmic Efficiency

Although all of Recut’s stages are fully parallel, Figure 5 illustrates the sequential runtimes of Recut and APP2 which to our knowledge has the current fastest reconstruction and skeletonization stages. In order to do an exact comparison, we had to run on vertex counts that APP2 can complete in a reasonable time-frame. It is common practice to run APP2 with a time-out since it has run-away runtimes on larger vertex counts.

**Fig. 5.**
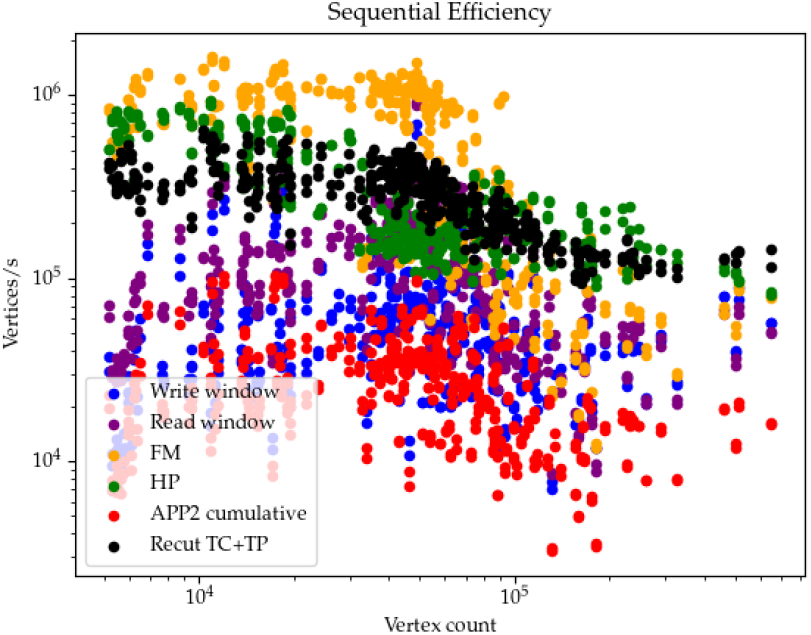
Sequential runtimes of APP2 and Recut stages plotted against the total voxel counts.

**Fig. 6.**
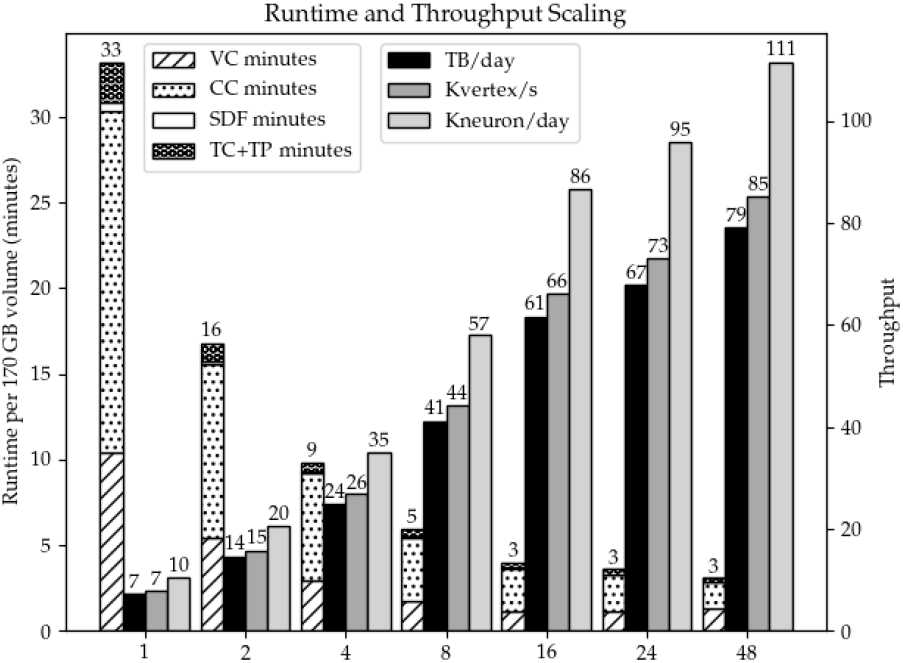
Runtime and throughput of Recut’s stages.

The throughput in Figure 5 also accounts for the steps APP2 must run without the Recut framework. The bounding box window must be written, read again by APP2 before the FM and HP stage. We do not include the time for a APP2 framework to read the entire image initially since this is usually semi-manual. In sequential settings, Recut’s prune stages are about an order of magnitude faster than APP2 in denser data where multiple trees are within a single connected component. Recut’s TC+TP stages have similar efficiency profiles to APP2’s HP, owing to the fact that they share the same algorithmic efficiency as shown in I.

### D. Peak Performance

The performance is even better than the algorithmic efficiency would suggest since n undergoes a substantial narrowing at the CC and TC stage as shown in Figure 1 and the downstream stages only visit the new sparse active sets. All Recut stages have a parallelized implementation, thus the *n* operations at each stage are conducted in parallel across available cores of a single CPU. These algorithmic efficiency improvements combined with the concurrent execution model translate to lower runtimes and higher pipeline throughput as illustrated in Tables IV and V. We also achieve lower runtimes on 16× larger images than the manually managed APP2.

**TABLE IV.** Recut runtimes per stage on confocal half coronal section with 30× objective lens on Camk-MORF3, 24-cores.

| Stage | Recut (s) |
| --- | --- |
| VC | 81.2 |
| CC | 91.1 |
| SDF | 1.95 |
| TC+TP | 11.6 |
| Hemi-section total | 186 |
| Striatum total | 2970 |

**TABLE V.**
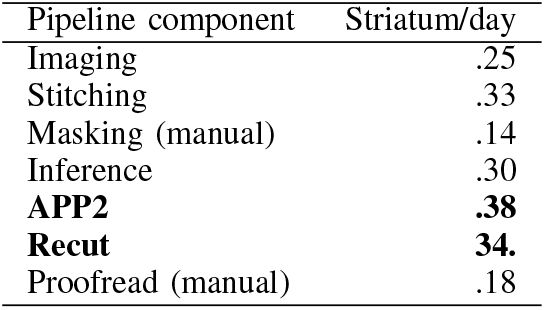
APP2 vs. Recut throughput per striatum brain region 30× objective lens, Camk-MORF3, 24-cores.

### E. Resource Utilization

The graph topology footprints are shown at the vertex scale in Tables II and III and at the macro scale in Table VI. To our knowledge, the 509× compression factor of an image via VDB is the largest compression factor found in the neuroscience community due to being uniquely suited to sparse volumes. To remain responsive at interactive time scales, Recut must have adequate system memory (DRAM) such that the full runtime footprint can fit. For the data sizes used in Table VI, this equates to about 6 GB, which easily fits in the memory of most modern laptops. This replaces the read and write steps of APP2 recorded in 5 with the VC stage on the whole image.

**TABLE VI.**
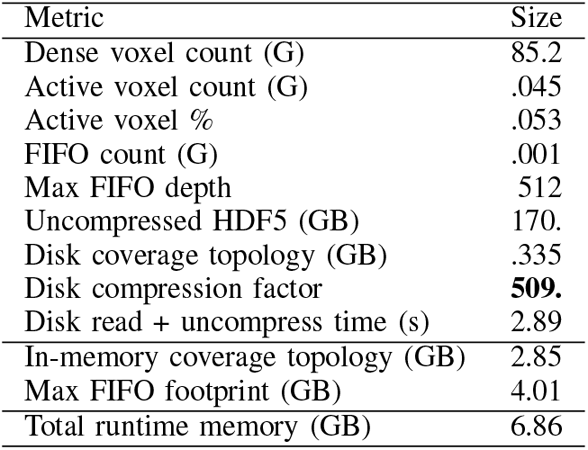
Data footprints for the striatum region of 1 hemi-section, 30× objective lens, Camk2a; MORF3 ~1% fluorescent labeling density, bounding volume of 10k,18k,468.

| Metric | Size |
| --- | --- |
| Dense voxel count (G) | 85.2 |
| Active voxel count (G) | .045 |
| Active voxel % | .053 |
| FIFO count (G) | .001 |
| Max FIFO depth | 512 |
| Uncompressed HDF5 (GB) | 170. |
| Disk coverage topology (GB) | .335 |
| Disk compression factor | **509.** |
| Disk read + uncompress time (s) | 2.89 |
| In-memory coverage topology (GB) | 2.85 |
| Max FIFO footprint (GB) | 4.01 |
| Total runtime memory (GB) | 6.86 |

### F. Scalability

While Recut succeeds in decoupling image IO from graph computation in reconstruction algorithms, it fails to achieve linear scaling on reconstructible data. This is due to the extreme sparsity of the data and the current parallelization strategy, as evidenced by the fact that a highly similar block-based fastmarching algorithm achieved super-linear speedup on dense volumes[6]. Both the SDF and CC stage apply the same paralellization strategy and traversal on the same set of nodes, yet SDF is 41× faster due to retaining starting seed points at all surface voxels. CC must start from single soma locations, which slows the propagation of the wavefront and dramatically increases the iterations until convergence.

On synthetically generated data where ground truth is known, 10% density is about 17× faster when using 16 cores. However, 10% pixel label density is beyond the upper limit for what is reconstructible on real data either for existing automated algorithms or for manual human reconstructions. At labeling densities above 1%, images are too saturated for proofreaders to effectively correct tree structures.

## V. Conclusions and Future Work

Lossless high resolution imaging can be less costly to store or transfer than even heavily downsampled images. While Recut is currently only supported for common PCs and CPUs, by comparison the average DRAM in newly purchased phones is almost 5 GB according to the Mobile Handset Sell-through Tracker. This work could enable productive and interactive workflows at the brain region vs. the single neuron scale on more common devices. However, to truly take advantage of these compression advantages at scale, a GUI tool for neuron proofreading should support visualization of VDB grids.

At a project scale, the accurately reconstructed neurons per animal–the *yield*–is a fundamental determinant of throughput. Yield is proportional to labeling density. Recut achieves consistent runtimes proportional to active voxel counts (instead of bounding volume) without hard to predict timeouts. Effectively leveraging sparsity in this way is the key to unprecedented image bounding volume scales.

We can also have predictable tradeoffs between label density and proofreadability, thus placing the reconstruction bottleneck back on yield per animal as opposed to image bounding volumes. Unified designs and simple implementations allow greater automation with less software infrastructure.

Recut makes large advancements in producing SWCs, the most common encoding of morphology today. However, richer or more native encodings such as the output windows may be more fruitful in combination with future NN-based approaches for morphological analysis. Recut is already particularly suited for efficiently generating training data for NN tools which suffer in this regard. Recut’s output windows can come from multiple image channels while allowing window enhancements, filtering, and positioning around neurons of interest.

Recut also assumes proofreading. This keeps humans in the loop for improving the neuron outputs before analysis. More aggressive and sophisticated filtering based on morphological metrics could mitigate or remove the need proofreading in certain scenarios. This would allow scientists to better leverage the yield and throughput benefits of this pipeline.

This work aims to place reconstruction automation, hierarchical parallel infrastructure and data-oriented programming as central concerns in the design of future neuroscience pipelines. Given the right abstraction, we have demonstrated that a high performance and scalable framework is possible even in unique and challenging data settings. Though beyond the scope of this report, this framework is modular and extends to several graphbased neural kernels in our own pipeline, providing similar performance and scalability gains. Having correctness, accuracy and performance isolated and tested automatically allows Recut to enforce established standards and prevents regressions on these key principles by new external collaborators. Through simple design, we can provide a hardware-agnostic scientific community an open source framework for performance, thus advancing the possibility of making real-time analysis or interactive data science frameworks the common workflow.

## A. Acknowledgments

We would like to thank the following authors for their contributions to this work. MZ, CC, HD, CP, WY, YC, ZC, and JC advised on design and provided helpful discussions and feedback. CP, CC and WY also provided the mouse imaging data. CC led the reconstruction proofread efforts. MZ designed and implemented the backend image IO library and the original sequential SG, CC, GC, and windowing output stages of the pipeline. KM designed, implemented and analyzed the Recut parallel framework and wrote the manuscript. We also thank Tony Nowatski for his suggestions on an early draft of the paper. This research is funded under NIH Grant No.: (U01MH117079-01).

## Notes

### Competing Interest Statement

The authors have declared no competing interest.

